# Ab-initio determination of the shape of membrane proteins in a nanodisc

**DOI:** 10.1101/2020.09.11.293043

**Authors:** Simone Orioli, Carl G. Henning Hansen, Lise Arleth

## Abstract

We introduce a new software, called *Marbles*, that employs SAXS intensities to predict the shape of membrane proteins embedded into membrane nanodiscs. To gain computational speed and efficient convergence, the strategy is based on a hybrid approach that allows one to account for the nanodisc contribution to the SAXS intensity through a semi-analytical model, while the embedded membrane protein is treated as set of beads, similarly to well known ab-initio methods. The code, implemented in C++ with a Python user interface, provides a good performance and includes the possibility to systematically treat unstructured domains. We prove the reliability and flexibility of our approach by benchmarking the code on a toy model and two proteins of very different geometry and size.

## 1. Introduction

Membrane proteins (MPs) constitute approximately 30% of the human proteome (Wallin & Heijne, 1998; Goffeau *et al.*, 1996) and are targets for approximately 60% of medical drugs (Bakheet & Doig, 2009; Yildirim *et al.*, 2007), but, among all the protein structures deposited in the Protein Data Bank, they represent only less than 3% of the total (Fagerberg *et al.*, 2010). This apparent paradox is explained by the technical difficulties hampering the crystallography-based experimental determination of their three-dimensional structures (Seddon *et al.*, 2004; Kang & Li, 2011; Rawlings, 2016). More recently, Cryo-Electron Microscopy (CryoEM) has undergone a so-called resolution revolution allowing for the visualisation of biomolecules, including membrane proteins, directly in 3D and with down to 2 Å resolution (Callaway, 2015). Still, however, there are several types of MPs that remain challenging also for CryoEM due to their too small size and/or their large flexibility.

These complications can be softened by the use of small angle x-ray scattering (SAXS) (Skar-Gislinge et al., 2010; Berthaud et al., 2012; Kynde et al., 2014; Dadinova et al., 2020), which, despite that it only provides a resolution down to about 10 Å, allows one to study complex biological samples directly in solution. Moreover, in recent years, nanodiscs (NDs), i.e. discoidal lipid bilayers enclosed by helical and amphipatic membrane scaffold proteins (Bayburt et al., 2002; Denisov et al., 2004; Denisov & Sligar, 2017; Hagn et al., 2013; Bibow et al., 2017; Johansen et al., 2019), have been established as a powerful tool to mimic native conditions for MPs. It has also been shown that small angle scattering (SAS) techniques can be systematically applied to study them (Skar-Gislinge et al., 2010; Kynde et al., 2014). As a part of this, new tools are being developed that allow for direct refinement of molecular models, generated from simulation, against the small-angle scattering data (Kassem et al., 2020; Bengtsen et al., 2020). This combination of a physically underpinned models and experimental data contains a huge potential for understanding the relevant solution properties of MPs and other biological macromolecules.

Despite their great versatility, SAS methods, when used on their own, have long been considered as techniques carrying a low information content, especially because of their resolution. This is a well known limitation induced by the signal-to-noise ratio in combination with orientational average employed to determine the one-dimensional scattering intensity. However, these techniques are in fact rich in information about the size and shape of biological samples (Jacques & Trewhella, 2010), as confirmed by Shannon-theory based methods to determine the information contents in SAS signals (Frieden, 1971; Vestergaard & Hansen, 2006). Indeed it is possible to deduce quite accurately the size and shape of a protein just from the knowledge of the corresponding SAXS intensity and by applying indirect methods for deducing the 3D shape of the underlying particles through simple physics-based constraints. The widely applied so-called ab-initio methods (Svergun, 1999; Svergun et al., 2001) provide an excellent example of this.

Relevant examples are present in the literature of MP shape determination in the case of proteins embedded in detergents based on SAXS (Pérez & Koutsioubas, 2015), contrast variation SANS at the match-out conditions for the detergents (Oliver et al., 2017), or SANS in combination with so-called stealth detergents (Midtgaard et al., 2018) or analogous stealth nanodiscs (Josts et al., 2018; Maric et al., 2014). In the case of SANS, the carrier system of the MP (e.g. the nanodisc) can be matched out by contrast variation, and this in principle allows one to determine the structure of the MP just by employing methods that are already available in the literature. However, high-intensity X-ray sources are much more abundant than neutron ones and, as a consequence, SAXS data are both more easily accessible and provide a higher information content than SANS (Pedersen et al., 2014) for solution based biomolecular systems. For this reason, it is relevant to develop approaches for interpreting SAXS data of MPs embedded in nanodiscs.

In this paper we try and overcome some of the current limitations of ab-initio methods by exploiting and extending a recently published hybrid approach (Kynde et al., 2014; Skar-Gislinge et al., 2015) and making it available through a novel software, called *Marbles*. The code implements an analytical model for the nanodiscs and treats the membrane protein as a set of beads, whose positions are optimized through the minimization of an empirical penalty function. The pipeline of the shape reconstruction is summarized in Fig 1: first, the information about the nanodisc is encoded in a set of parameters by fitting an analytical model to the SAXS signal of a solution of empty nanodiscs; second, both these fit parameters and the SAXS signal of a solution of nanodisc-bound MPs are fed to *Marbles*, which employs them to minimize a penalty function and consequently recover the shape of the protein of interest.

**Fig. 1.**
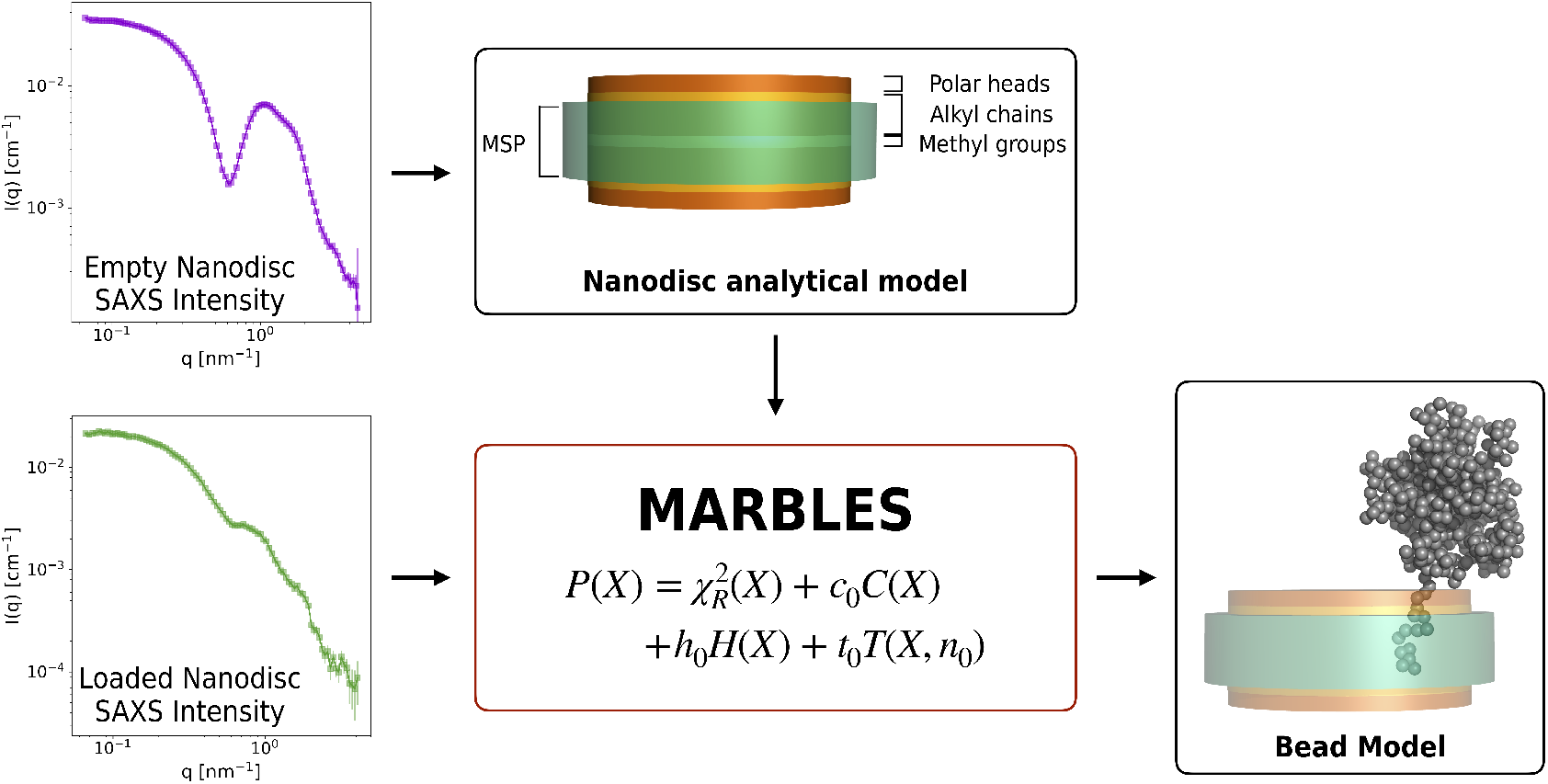
Schematic representation of the pipeline employed to recover the shape of a membrane protein in a nanodisc using *Marbles*.

*Marbles* is based on a previously published work (Skar-Gislinge et al., 2015), which was optimized to solve the problem for a single dataset on a specific member of the Cytochrome P450 family (more details in section 4.1). The software we introduce in this manuscript, instead, extends its capabilities by making it readily applicable to any protein of interest, including membrane proteins with unstructured domains. *Marbles* features a new simple strategy for the determination of the position of the starting configuration that substantially accelerates convergence. This, together with a complete restyling and optimization of the code and a better assessment of some hyperparameters allows us to get a 10-fold speed up with respect to the previous version, making *Marbles* a valuable tool for in-situ analysis of experimental data.

The article is structured as follows. First, we briefly discuss the mathematical model employed in the code (section 2), the algorithm applied to retrieve the protein structure and the code implementation (section 3). We test *Marbles* on a toy model (appendix B) and on two test proteins (section 4), cytochrome P450 3A4 and tissue factor, and we show how the results are consistent with previous knowledge on the systems in the literature. Finally, we discuss some relevant points on how to interpret the results and how the code can be possibly extended (section 5) and we close with some concluding remarks (section 6).

## 2. Physical model

Let us start from the well known fact that the scattering amplitude A(**q**) of a system can be expressed via an expansion on the basis of spherical harmonics *Y_lm_*(Ω) (Svergun *et al.*, 1995):

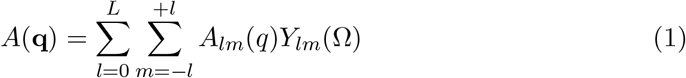

where *L* is the maximum number of basis elements employed in the expansion, i.e. the quantum angular momentum and *m* is a generalized magnetic quantum number, while *A_lm_*(*q*) are the (complex) partial amplitudes and depend on the shape and on the excess scattering length Δ*b* of the system under study (e.g. a protein). The computation of partial amplitudes requires an orientational average over all the solid angle Ω, while the angular dependence of the scattering amplitude is moved to the spherical harmonics. The corresponding scattering intensity is then obtained as

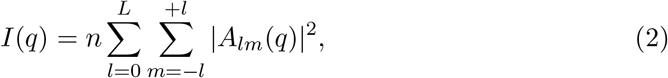

where *n* is the solute number density. Eq. 1 is a rather general expression and will serve as a starting point to build our hybrid model. Its construction will proceed as follows. In section 2.1 we will discuss how proteins can be treated as sets of beads; in section 2.2 we will present an analytical formulation of a membrane nanodisc form factor and, finally, in section 2.3 we will show how to incorporate the two descriptions into a hybrid approach.

### 2.1. Proteins

Given the relatively low resolution of SAS techniques (Jacques & Trewhella, 2010) and in order to decrease the complexity of the problem, it is customary to coarse grain protein amino acids into material points, or *beads*. Within this simple assumption, the partial amplitudes in Eq. 1 can be computed analytically (Svergun *et al.*, 1995):

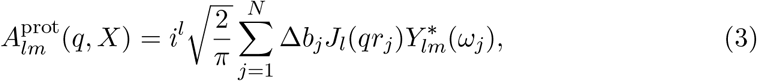

where *X* = (**r**_1_,…, ***r***_*N*_) is the protein’s configuration, *i* is the imaginary unit, *N* is the number of beads, Δ*b_j_* is the excess scattering length of the *j*-th bead, *J_l_*(*qr_j_*) is the *l*-th Bessel function of the first kind, 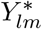 is the complex conjugate spherical harmonics and the position of the *j*-th bead in real space is given by **r**_*j*_ = (**r**_j_,*ω_j_*). Eq. 3 is an especially handy expression, as it displays a feature which is crucial in applications such as ab-initio bead modeling. Indeed, if the *i*-th bead is moved from it’s original position **r**_*j*_ to a new location **r**_*j*_ + *δ***r**_*j*_, the expansion coefficient *A_lm_*(*q*) of the new configuration can be obtained by subtracting the contribution of the bead in **r**_*j*_ and summing the new intensity component in **r**_*j*_ + *δ***r**_*j*_. This is a rather computationally inexpensive task, as it comes without the need to re-compute the contributions of all the other beads.

### 2.2. Nanodiscs

As described in previous works (Skar-Gislinge *et al.*, 2010; Skar-Gislinge & Arleth, 2011), the form factor of a nanodisc can be approximated by a combination of the form factors of suitably stacked cylinders with elliptic cross-section. Each cylinder represents different structural features of the nanodisc, in particular (see the upper central picture in Fig. 1): i) the top and bottom layers are employed to represent the lipid heads; ii) two further layers are used to describe the lipid alkyl chains; iii) the innermost layer represents the lipid methyl groups. A further hollow cylinder is added around this construct, and it is used to describe the membrane scaffold protein (MSP or protein belt) surrounding the lipid bilayer. We stress here that the methyl layer is added for consistency with previously published works (Skar-Gislinge *et al.*, 2010; Skar-Gislinge & Arleth, 2011; Skar-Gislinge *et al.*, 2015) but it is not strictly required and might be easily removed.

This modeling strategy is particularly convenient because of the additive nature of form factor amplitudes and because the form factor for a cylinder is known semi-analytically (Mittelbach & Porod, 1961; Pedersen, 1997). Therefore, by simple geometrical arguments it is possible to deduce the form factors of all the nanodisc components. The sum of their contributions, weighted with the corresponding average excess scattering lengths, provides the overall scattering form factor amplitude of the nanodisc, *A_nd_*:

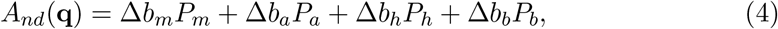

where *P_m_, P_a_, P_h_* and *P_b_* are the form factor amplitudes of, respectively, the methyl groups, alkyl chains, head groups and belt protein layers multiplied by the corresponding excess scattering lengths Δ*b*, which, as usual, are calculated based on molecular constraints (see more details on this in section 3.1). As also discussed in previous works (Skar-Gislinge & Arleth, 2011), however, this model does not account for the interface roughness observed in scattering experiments on real samples. The infinitely sharp interfaces of the cylinders in the mathematical model result in artifacts in the high-*q* region of the scattering intensity. To correct for this behavior, a Gaussian roughness term *e*^−(*qR*)2^ is multiplied to the nanodisc scattering amplitude, where *R* measures the roughness of the nanodisc surface (Als-Nielsen & McMorrow, 2011).

### 2.3. Hybrid model: Membrane protein in a nanodisc

To treat the nanodisc and the beads intensities on the same footing, we can expand *A_nd_*(**q**) too in the basis of spherical harmonics. Given the complexity of the model employed to describe the nanodisc, however, analytical integration of the orientational average in the partial amplitudes is out of question. The calculation can be therefore discretized on a 2*L* × 2*L* grid, leading to (Kynde *et al.*, 2014; Mohlenkamp, 1999; Mohlenkamp, 2016):

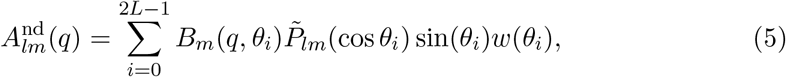

where 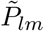 are the L2-normalized Legendre polynomials, while

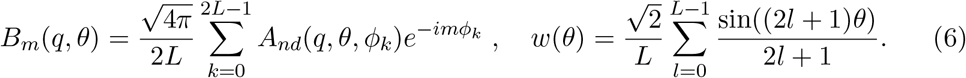

A key note is in place here. Because we are dealing with membrane proteins, it is clear that part of the protein, described through eq. 3, will be embedded in the nanodisc. Therefore, we distinguish among the four different backgrounds in which beads can be immersed in: solvent, lipid polar heads, lipid alkyl chains or lipid methyl groups. Depending on the location of the bead in the disc and hence its background, the excess scattering length of the beads, Δ*b_j_* of eq. 3 will change accordingly (Kynde et al., 2014; Skar-Gislinge et al., 2015).

### 2.4. Membrane protein with unstructured domain

The strategy presented in section 2.3 is sufficiently general to allow for further refinements of the model, when required by complex samples. In particular, as we will see in section 4.2 with the application on tissue factor, one might need to account for the presence of disordered regions of the polypeptide chain protruding from the bottom leaflet of the lipid bilayer, e.g. a histidine tag or an intrisincally disordered intracellular domain (Bugge et al., 2016). We consider such unstructured region to be well described by a Gaussian random coil (Debye, 1947; Hammouda, 1992; Pedersen & Gerstenberg, 1996) attached to the bottom leaflet of the nanodisc. The orientationally avaraged scattering amplitude of such an object is given by the Hammouda term (Hammouda, 1992):

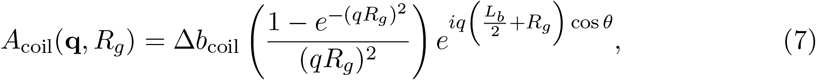

where the first bracket is the orientationally averaged scattering amplitude of the coil, *R_g_* is the radius of gyration of the unstructured coil, *L_b_* is the height of the nanodisc bilayer and the exponential phase factor is needed to displace the coil on the bottom leaflet of the nanodisc along the *z*-axis, normal to the nanodisc plane. The radius of gyration can either be taken as a fit parameter or it can be estimated through the empirical scaling law determined by Kohn and collaborators (Kohn et al., 2004), *R_g_* = *R_0_N^ν^*, where *R*_0_ ~ 1.927Å, *N* is the number of residues composing the random coil and *v* = 0.598 is a scaling constant. To make Eq. (7) compatible with our hybrid model, we can also expand *A*_coil_(**q**) on the basis of spherical harmonics. The expansion coefficients 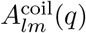 can be obtained following Eq. 5 for the case of the nanodisc. The last step in building the model requires to note that the coil self-correlation term in the scattering intensity is not well described by 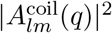, but rather it has to be treated separately by a Debye term (Debye, 1947; Pedersen & Gerstenberg, 1996):

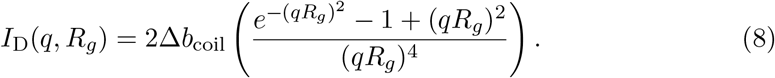

The overall scattering intensity of the hybrid model is then provided by

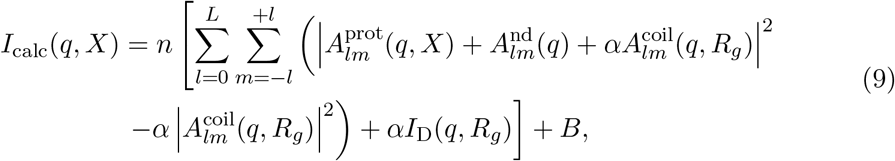

where *n* is the particle number density and *B* is a constant background, which should be ideally very close to zero. Setting *α* = 0 is equivalent to disregard the intensity components coming from a random coil, while *α* = 1 includes them. The normalization provided by the particle number density sets the intensity to absolute scale, but when experimental data are not provided in such a normalization (which is the case of the results discussed in section 4), one can rescale the calculated intensity by a factor *I*_exp_(0), which can be determined either by a Guinier fit or by indirect Fourier transform of the experimental SAXS intensity.

## 3. The Algorithm

The physical description of a membrane protein in a nanodisc we summarized in Eq. 9 provides a simple forward model that can be applied to compute their scattering intensity. In this section we will make use of this forward model to design an algorithm able to retrieve the shape of a membrane protein in an ab-initio fashion. To do so, we take inspiration from the implementation of GASBOR (Svergun *et al.*, 2001) and later works (Franke & Svergun, 2009; Skar-Gislinge *et al.*, 2015), where the shape of a protein is determined by a Monte Carlo procedure which minimizes the discrepancy between the experimental scattering intensity and the one computed from a set of dummy beads. Here we follow a similar philosophy and we further extend the code to allow for the description of nanodiscs.

### 3.1. Retrieval of Nanodisc Parameters

The first step in the determination of the membrane protein shape is the fit of the nanodisc model in section 2.2 to the scattering intensity obtained from a solution of empty (or unloaded) nanodiscs. Indeed, if the protein is not substantially perturbing the nanodisc (see the discussion on this assumption in section 5.1), this strategy provides the simplest way to determine the parameters describing the nanodisc model.

In practice, the fit is carried out using the *WillItFit* software (Pedersen et al., 2013). Despite the relative complexity of the nanodisc model, which allows in principle for a large set of free parameters, *WillItFit’*s approach of introducing molecular constraints to fix the interdependence between chemico-physical quantities (Skar-Gislinge et al., 2010; Skar-Gislinge & Arleth, 2011) reduces the number of free parameters to 8: the nanodisc axis ratio *ε*, the number of lipids per leaflet *N_l_*, the average area per lipid headgroup *A_h_*, the height of the MSP *L_b_*, the surface roughness R, the constant background *B* and the correction factors to the volumes of MSP and lipids, respectively *C_b_* and *C_l_*. While the first six parameters have a clear physical interpretation, the last two are used to allow some additional freedom in the model by slightly perturbing the expected molecular volumes of lipids and the belt protein, and hence their excess scattering length densities. Both of them are multiplicative factors and deviations from unity are expected to be no more than 10%. Other physical features of the nanodisc, e.g. the heights of the different nanodisc layers, can be deduced a posteriori from just these 8 parameters, the relationships connecting them and the chemical knowledge about the MSP and the lipids composing the bilayer (Skar-Gislinge et al., 2010). Furthermore, the height of the MSP, *L_b_* is often fixed to 25.8Å, as determined by NMR (Bibow *et al.*, 2017; Johansen *et al.*, 2019).

### 3.2. Simulated Annealing

Once the nanodisc parameters have been determined, a set of beads is used to retrieve the shape of the membrane protein inserted in the bilayer. In particular, we seek for the three-dimensional disposition of beads in the nanodisc whose scattering intensity minimizes the discrepancy with the experimental one. A measure of this discrepancy is naturally provided by the reduced chi-squared:

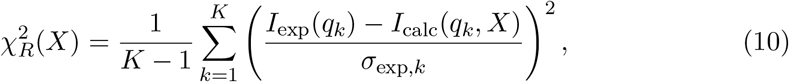

where *K* is the number of intensity points. However, the ensemble of all the possible configurations of *N* beads that minimize 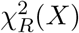 is extremely large and, among all these, only an infinitesimal fraction resembles a protein shape. For this reason, further constraints need to be added to restrict the search among bead assemblies that are protein like. The definition of what is a protein like shape is somewhat arbitrary, but minimal requirements ask for the protein configuration to be connected and to follow the typical nearest-neighbour distribution of residues in a protein (Svergun et al., 2001). In addition, a term that enforces the embedding of part of the protein and the nanodisc can be added (Skar-Gislinge et al., 2015). These features can be enforced by adding them as further constraints, together with the 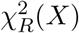, so to define an overall penalty function:

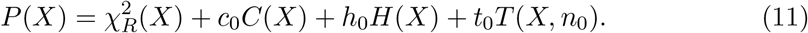

*C*(*X*) is the component of the penalty function that enforces the set of beads to be interconnected, and it does it by comparing the length, in number of beads, of the longest graph that can be built from the current configuration, *N*G**, with the total number of beads in the model *N*, *C*(*X*) = log(*N_G_/N*). *H*(*X*) is the portion of the penalty ensuring a protein like nearest-neighbour distribution of beads. This is done by comparing the distribution obtained from configuration *X* with an average one, determined from a set of proteins in the Protein Data Bank (Svergun et al., 2001), using a *χ*^2^ function. Finally, the last term enforces the interaction between the protein and the nanodisc by rewarding the insertion of *n*_0_ beads into the bilayer, *T*(*X, n*_0_) = (*n*(*X*) – *n*_0_)^2^, where *n*(*X*) is the number of beads embedded in the nanodisc for configuration *X*.

The optimal three-dimensional disposition of beads is realized when *P*(*X*) is minimal. To look for such minimum, we employ simulated annealing (Kirkpatrick et al., 1983) in the way outlined in the following (see also Fig. 2):

i. *N* beads, where *N* is the number of protein residues, are randomly placed within a sphere of diameter *D*, positioned at an altitude *z*_0_ over the nanodisc center. The initial intensity is then estimated using Eq. 9;
ii. the position of the sphere is optimized by running a grid search over many *xy* translations within the smallest rectangle confining the nanodisc. Because of the two-fold symmetry of the nanodisc, the search is carried out only in the upper right corner of the ellipse. The optimal location of the initial sphere is selected as the one minimising the 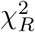 between the computed and the experimental SAXS intensity. The sphere is then translated to the optimal position, defining an initial configuration *X*_0_;
iii. two beads *j* and *k* are drawn randomly from the set, and *j* is moved in the vicinity of *k* along a random vector **r**_move_ of norm 3.8 Å (the typical distance between *C_α_* in a protein, (Chakraborty et al., 2013)). If *j* is found to be closer than a distance *r*_clash_ = 1.8Å to any bead in the set, the move is rejected and a different random vector is generated. This last operation is repeated up to 1000 times, and if all the moving attempts are rejected a new pair of beads is drawn. Depending on the new position of the bead and the corresponding background in which the bead is embedded, its corresponding excess scattering length is evaluated;
iv. if the movement of bead *j* is not rejected, the system describes a new configuration *X*_1_ and the value of the penalty function *P*(*X*_1_) is evaluated. Following a Metropolis criterion (Metropolis et al., 1953), the new configuration is accepted with a probability *p*(*β*_0_,*X*_0_,*X*_1_) = min[1,exp{−*β*_0_(*P*(*X*_1_) – *P*(*X*_0_))}], where *β*_0_ is the initial effective inverse temperature. This means that every time a configuration is evaluated, a uniform random number *ρ* ∈ [0,1] is drawn. If *p*(*β*_0_,*X*_0_,*X*_1_) > *ρ* the move is accepted, otherwise it is rejected and the original system’s configuration *X*_0_ is restored;
v. steps (iv) and (v) are repeated until *N* moves have been accepted. At that point, a cooling schedule is applied and the effective temperature 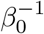 is decreased by multiplying it by the cooling rate *s* < 1, 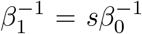. After this operation, we say a pass has been completed;
vi. step (vi) is iterated until a convergence criterion is reached. In particular, a lower threshold is set for the effective temperature, 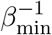, and the simulation is stopped at pass k if 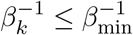.

**Fig. 2.**
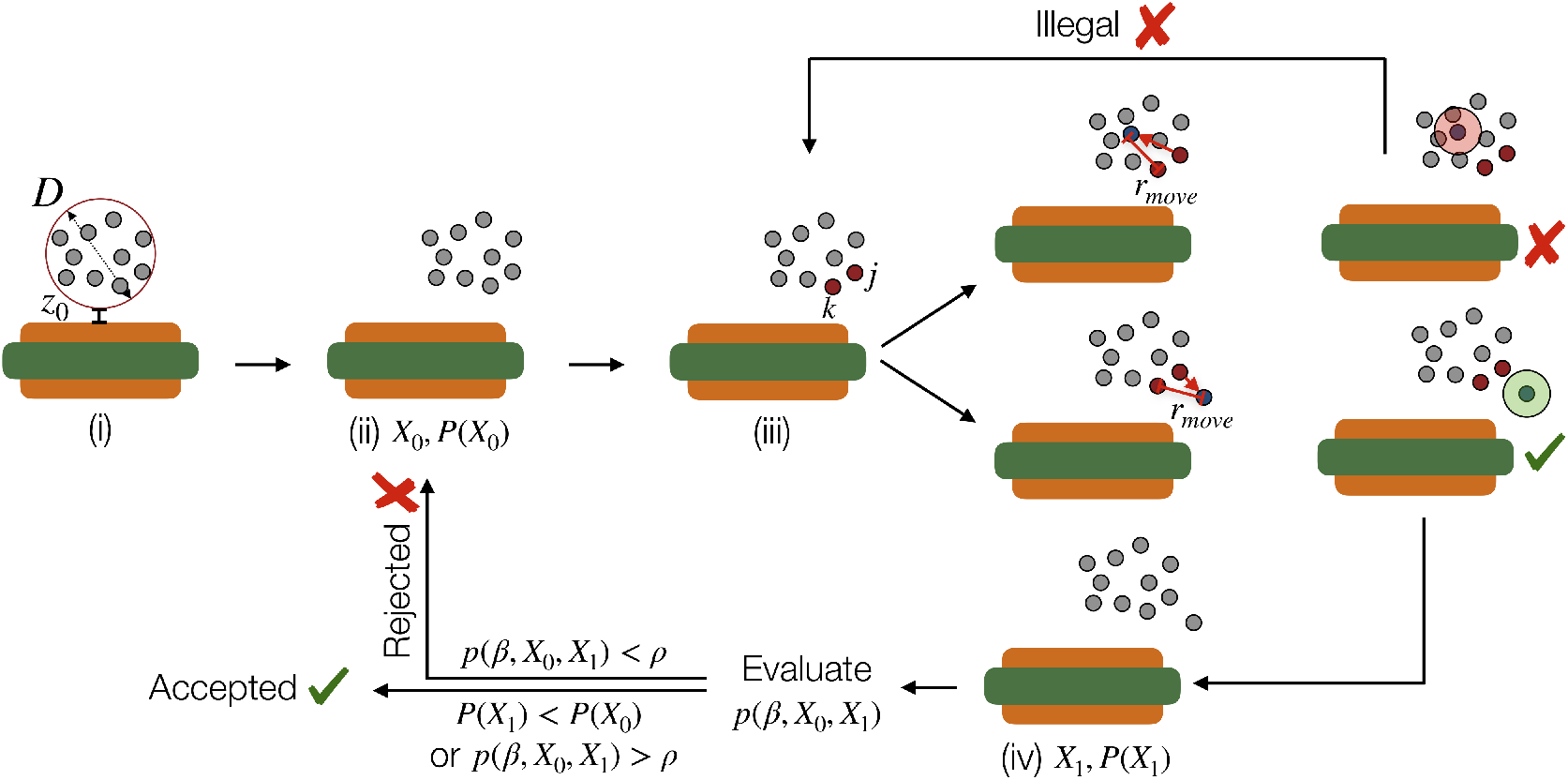
Graphical summary of a simulated annealing step. The roman numbers in brackets are used to refer to the different stages of the algorithm.

The final configuration *X*_final_ is interpreted as the best estimation of the membrane protein shape. Because of the under-determined and stochastic nature of this approach, it is desirable to run the optimization multiple times with the same data. This has multiple advantages: first of all, it allows to determine if some runs ended up stuck in a local minimum; secondary, it makes the shape estimation more robust via the exclusion of outlier beads by averaging over multiple final conformations. The latter procedure can be carried out, for example, using DAMAVER (Volkov & Svergun, 2003).

Point (ii) of our simulated annealing algorithm deserves some additional comments. Indeed, the initial search for the optimal position of the initial sphere of beads is carried out in order to make the optimization more efficient by starting it from a more favourable local minimum in the penalty landscape. To show that this procedure is able to determine a reasonable starting position on the nanodisc, we generated synthetic SAXS data by embedding a protein (cytochrome P450 3A4, more details on the protein in section 4.1 and on the generation of synthetic data in appendix A) in the middle of the nanodisc and close to the nanodisc’s rim. As discussed in detail in appendix B, the analysis of these toy models led us to the conclusion that if information about protein’s location on the nanodisc is encoded in the SAXS data, then it is possible to systematically take it into account to accelerate convergence by performing the grid search described in (ii).

### 3.3. Implementation and performance

We implemented *Marbles* in C++11 featuring a Python layer (both a parser and a GUI) for user’s simplicity of use. The Python component of the code is only concerned with the interpretation of the user’s input, while the whole calculation is carried out by the C++ layer. The code is freely available at https://github.com/Niels-Bohr-Institute-XNS-StructBiophys/Marbles, together with a complete user guide and examples of application. The code is released under the GPL-3.0 license, so it is free to be modified and redistributed within the conditions of the aforementioned license.

A single *Marbles* run of the two systems described in section 4 required less than 5 minutes to converge on a 2.3 GHz Intel Core i5 processor. The performance is 10 times higher than the one of the previous code (see appendix C) and it is compatible with a high-throughput application of the code on a small work station, or for quick data analyses to be performed directly at the beamline. Additional details on the code are provided in appendix D.

## 4. Applications

### 4.1. Case 1: Cytochrome P450 3A4

Cytochromes P450 belong to the superfamily of haem enzymes (Nelson, 2009) that are essential proteins for biosynthesis and catabolism of xenobiotics (De Montellano, 2005). They are monotopic proteins and as such, the mode of incorporation of the protein in the membrane is critical for the correct function of cytochromes P450 (Grinkova et al., 2013). In this application, we focus on the case of cytochrome P450 3A4 (CYP3A4), which interacts with the membrane through a short, 15 residues long, α-helix. The mode of incorporation of human cytochromes P450 into the membrane has been the subject of numerous molecular dynamics studies (Otyepka et al., 2012; Denisov et al., 2012; Baylon et al., 2013) and was also studied using a previous version of the code presented here (Skar-Gislinge *et al.*, 2015). Given the wide amount of literature on the subject, this protein provides a good benchmark for *Marbles*’ predictive capabilities.

Here, we used some previously published SAXS data on CYP3A4 (Skar-Gislinge et al., 2015) and provide them as an input to *Marbles* to recover the shape of the protein in native conditions.

First of all, we fitted the analytical nanodisc model, discussed in section 2.2, to the SAXS signal measured from a solution of empty nanodiscs. The fit result is shown in Fig. 3 (a), where the SAXS intensity of the solution of empty nanodiscs is reported as blue dots and the corresponding fit is shown as a black line superimposed to the data. In the left-hand side of Tab. 1, we report the parameters of the nanodisc model determined from the fit, together with their corresponding uncertainties.

**Fig. 3.**
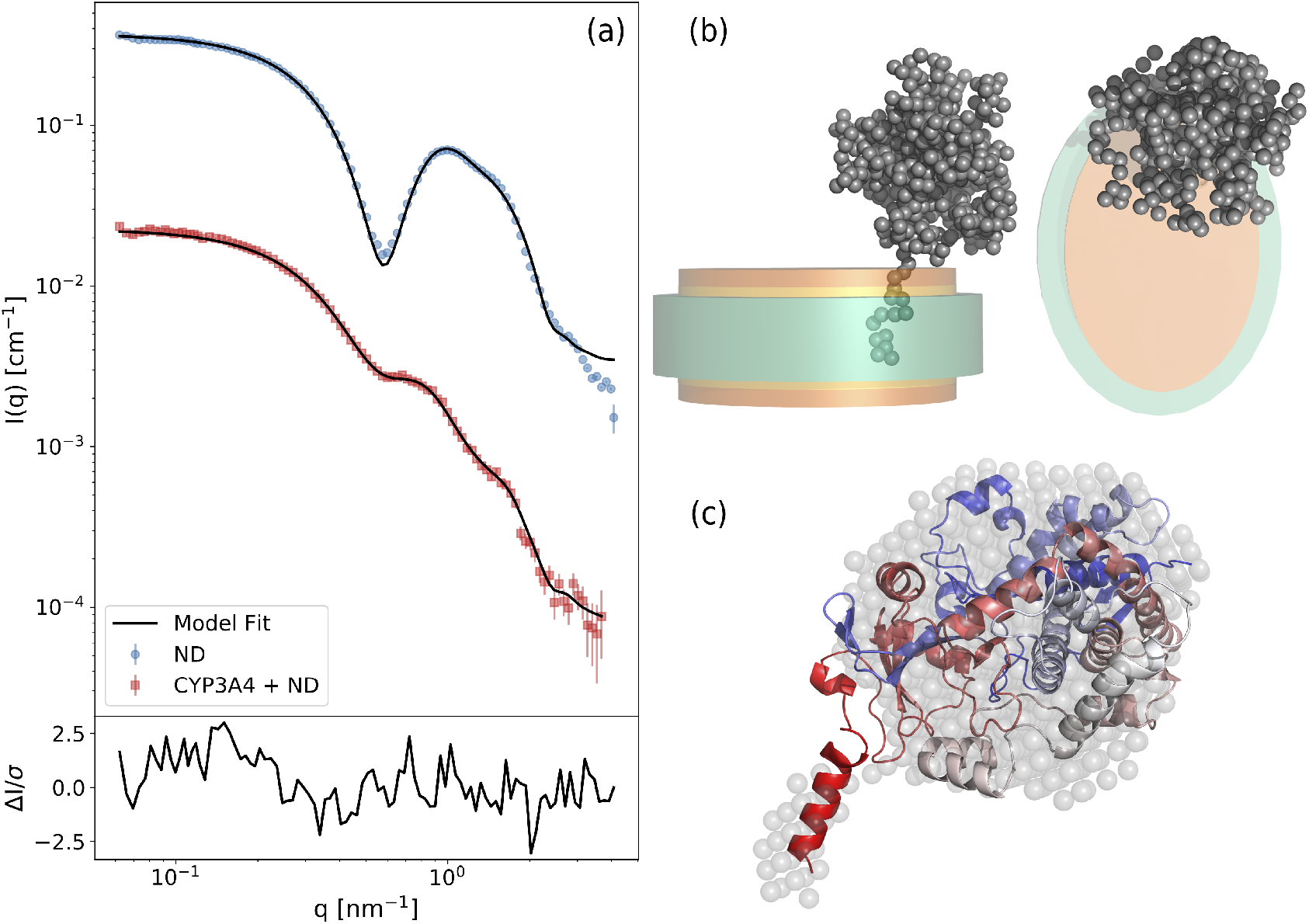
(a) SAXS data for the solution of empty nanodiscs (blue) and the solution of nanodiscs with reconstituted CYP3A4 (red). Experimental data are previously published in (Skar-Gislinge *et al.*, 2015). The full black curves show the corresponding model fits: the nanodisc model has been obtained with *WillItFit*, while the other one with *Marbles.* The lower panel reports the residuals between the experimental and the computed signal in the case of the reconstituted CYP3A4; (b) a typical configuration obtained by *Marbles* for CYP3A4 seen from the side (left) and from above (right); (c) superposition between the consensus model (transparent grey spheres), obtained by aligning 10 *Marbles* models using DAMAVER, and the CYP3A4 model obtained from MD simulations (cartoon).

**Table 1.** Right: parameters employed to fit the analytical nanodisc model to the SAXS intensity of a solution of empty nanodiscs. See section 3.1 for their meaning. Left: parameters employed in the 10 Marbles runs. See section 3 for their meaning.

| *parameters employed in the 10 Marbles runs. See section 3 for their meaning.* |  |  |  |
| --- | --- | --- | --- |
| Fit parameters | Values | Running parameters | Values |
| $\varepsilon$ | $1.45 \pm 0.05$ | $c_0$ | 300 |
| $N_l$ | $127 \pm 4$ | $h_0$ | 1 |
| $A_h$ ($\text{\AA}^2$) | $64.2 \pm 0.8$ | $t_0$ | 5 |
| $L_b$ ($\text{\AA}$) | 25.8 | $n_0$ | 15 |
| $R$ ($\text{\AA}$) | $4.6 \pm 0.1$ | $r_{\text{clash}}$ ($\text{\AA}$) | 1.8 |
| $B$ ($\text{cm}^{-1}$) | $-(3.4 \pm 0.5) \times 10^{-4}$ | $\beta_0^{-1}$ | $\chi_R^2(X_0)/10$ |
| $C_b$ | $0.906 \pm 0.003$ | $s$ | 0.9 |
| $C_l$ | $1.034 \pm 0.003$ | $\beta_{\text{min}}^{-1}$ | $5 \times 10^{-3}$ |

After the computation of the optimal nanodisc parameters, we performed 10 independent runs with *Marbles*. The parameters we employed for all the runs are reported in the right-hand side of Tab. 1. All the 10 runs reached the selected convergence temperature within 60 passes, and we manually inspected the corresponding final configurations to check that they were properly embedded in the nanodisc and connected. During the Monte Carlo runs, the penalty function decreased from an average value of 1100 to an average of approximately 1.52. A typical fit achieved by *Marbles* is shown as a black line in Fig. 3 (a), superimposed to the red squares representing the SAXS intensity of a solution of CYP3A4 embedded in a nanodisc. The bottom part of Fig. 3 (a) reports the residuals of the corresponding fit.

A characteristic protein configuration resulting from the optimization is shown in Fig. 3 (b). Marbles predicts a stalk-like shape for the protein, with a rod-shaped intramembrane domain accomodated in the nanodisc and a bulky extracellular domain. All the generated models suggest a strongly decentered CYP3A4, in close proximity of the rim of the nanodisc, in accordance with previous studies (Denisov et al., 2005; Skar-Gislinge et al., 2015).

To check the consistency of our results against previous Molecular Dynamics (MD) simulations of CYP3A4 (Denisov et al., 2012), we employed DAMAVER (Volkov & Svergun, 2003) to average our 10 configurations and generate a consensus model (see Fig. 3 (c)). Later, we aligned the resulting model to the MD one and computed the normalized spatial discrepancy (NSD) between the two structures, resulting in NSD = 0.94. Given that a NSD below 1 is usually considered as an indication of structural similarity (Svergun et al., 2001), we can conclude that the shape of our consensus model is realistic and comparable with the one predicted by MD simulations of CYP3A4.

### 4.2. Case 2: Tissue Factor

Tissue Factor (TF) is a protein playing a key role in the blood coagulation cascade (Mackman, 2009) and, like most of single-pass membrane proteins (Bugge et al., 2016), it is composed by three domains: a soluble extracellular domain (sTF), an alpha-helical transmembrane domain and a short, disordered intracellular domain. While the high-resolution structure of sTF has been known for more than two decades (Banner et al., 1996; Sen et al., 2009), the structure of the membrane-bound TF remains elusive.

In a recent work, Tidemand et al. (Tidemand et al., 2020) have shown how TF can be reconstituted into POPC-POPS nanodiscs based on csMSP1E3D1 belts, which are extended, circularized and solubility enhanced as compared to the original MSP1D1 nanodisc belts. Here, we apply *Marbles* to determine the shape of the Tissue Factor in a nanodisc, using the resulting and already published SEC-SAXS data (Tidemand et al., 2020).

Following the same strategy outlined in the cytochrome P450 application, section 4.1, as a first step we fitted the SAXS data of a solution of empty nanodiscs using *WillItFit* (see the black line superimposed to the blue dots in Fig. 4 (a)), obtaining a set of parameters that we have then employed in our ab-initio modelling approach. The list of fit parameters is shown in Tab. 2 on the left-hand side. Later, we performed 10 independent *Marbles* run to obtain as many independent models. In the original work (Tidemand *et al.*, 2020), TF protein was purified together with an intracellular histidine tag, i.e. a 6-histidines long unstructured chain used as a tag for protein purification. Since this portion of the protein will also provide a component of the overall scattering intensity, we included the corresponding contribution as outlined in section 2.4.

**Fig. 4.**
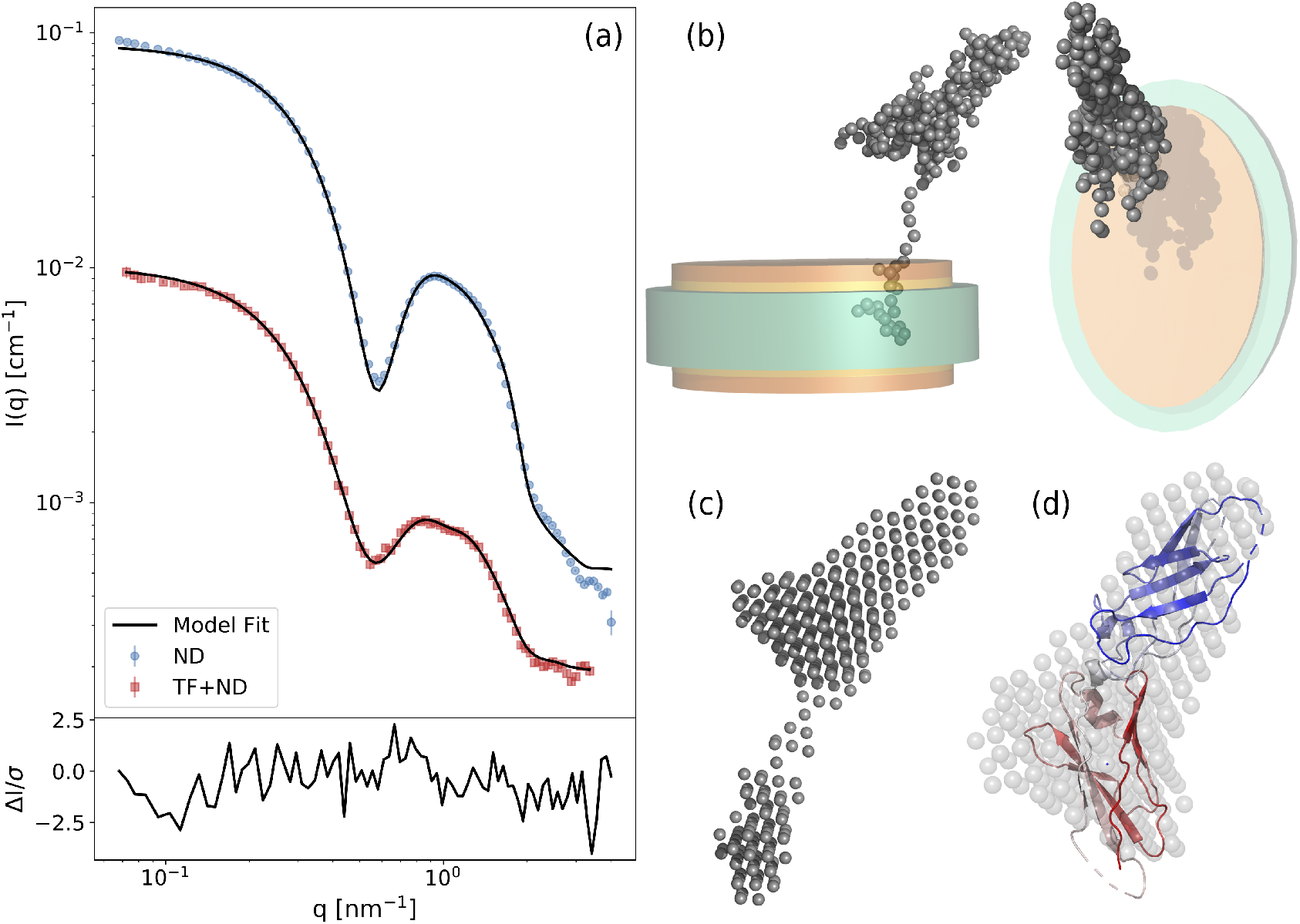
(a) SAXS data for the solution of empty nanodiscs (blue) and the solution of nanodiscs with reconstituted TF (red). The full black curves show the corresponding model fits: the nanodisc model has been obtained with *WillItFit*, while the other one with *Marbles*. The lower panel reports the residuals between the experimental and the computed signal in the case of the reconstituted TF; (b) a typical configuration obtained by *Marbles* for TF seen from the side (left) and from above (right); (c) consensus model obtained by aligning 10 *Marbles* models using DAMAVER; (d) superposition between the sTF crystal structure (cartoon) and the corresponding portion in the consensus model (transparent grey spheres).

**Table 2.** *Right: parameters employed to fit the analytical nanodisc model to the SAXS intensity of a solution of empty nanodiscs. See section 3.1 for their meaning. Left: parameters employed in the 10* Marbles *runs. See section 3 for their meaning*.

| *parameters employed in the 10 Marbles runs. See section 3 for their meaning.* |  |  |  |
| --- | --- | --- | --- |
| Fit parameters | Values | Running parameters | Values |
| $\varepsilon$ | $1.57 \pm 0.02$ | $c_0$ | 300 |
| $N_l$ | $175 \pm 2$ | $h_0$ | 1 |
| $A_h$ ($\text{\AA}^2$) | $58.1 \pm 0.5$ | $t_0$ | 5 |
| $L_b$ ($\text{\AA}$) | 25.8 | $n_0$ | 22 |
| $X$ ($\text{\AA}$) | $4.46 \pm 0.04$ | $r_{\text{clash}}$ ($\text{\AA}$) | 1.8 |
| $B$ ($\text{cm}^{-1}$) | $-(5.2 \pm 0.2) \times 10^{-4}$ | $\beta_0^{-1}$ | $\chi^2(X_0)/10$ |
| $C_b$ | $0.931 \pm 0.005$ | $s$ | 0.9 |
| $C_l$ | $1.009 \pm 0.002$ | $\beta_0^{\text{min}}$ | $10^{-3}$ |

The average value of the penalty function during the optimization performed by *Marbles* drops from 2200 to 1.78 in less than 80 passes. A typical fit is shown as a black line superimposed to the red squares in Fig. 4 (a), while a representative model is shown in Fig. 4 (b).

The resulting protein model displays an elongated shape, with a flexible linker connecting the sTF to the intramembrane helix. We want to stress that the results obtained on this protein strikingly differ from the ones on CYP3A4, as was initially expected because of their very different geometries. This prediction is conserved over all the models we generated, as confirmed by the consensus model calculated by aligning our ten models with DAMAVER. Fig. 4 (c) clearly shows an envelope which is coherent with the conformation of the extracellular domain sTF. This is confirmed by an *NSD* = 0.98 obtained aligning the crystal structure of the extracellular domain sTF (PDB code: 1TFH) with the corresponding motif in the consensus model (see Fig. 4 (d)). Further inspection of the consensus model reveals a feature resembling an elongated linker connecting the intramembrane domain with the sTF. The full TF construct indeed features a flexible linker, but in the literature not much is reported about its behavior. We argue that the predictions of our bead model are consistent with a dynamic behavior of the protein, even though it is hard to tell whether the measure of the linker’s extension is a true feature of the protein or a an artefact stemming from the fact that we are trying to fit a dynamical ensemble with a single structure (see 5.1). Our conclusion is that, overall, *Marbles* results are consistent with the available experimental knowledge on Tissue Factor.

## 5. Discussion

### 5.1. Fundamental assumptions

It is worth stressing here that the algorithm behind *Marbles* is designed to work under some well defined physical assumptions. First of all, we assume the nanodisc not to be substantially perturbed by the interaction with the protein. This premise allows us to describe the nanodisc using immutable parameters that will be fixed at the beginning of the optimization procedure and will not be modified by the relative orientations of protein beads. Clearly, a major drawback of this assumption resides in the fact that it might limit the applicability of the software to single pass membrane proteins, i.e. proteins interacting with the nanodisc only via a thin intra-membrane anchor. In principle, it is possible to allow for some flexibility in the nanodisc parameters, i.e. by defining some ad-hoc Monte Carlo moves within physically reasonable boundaries. However, this would come with a cost in terms of convergence time, as the additional freedom would increase the frustration of the energy landscape. For this reason, we believe that methods such as MEMPROT (Pérez & Koutsioubas, 2015), MONSA (Svergun, 1999) or the rigid body modeling implemented in *WillItFit* (Skar-Gislinge *et al.*, 2015) would be more suited for proteins strongly perturbing the membrane (see discussion in section 5.2).

A further assumption we make is that we can safely ignore the component of the intensity coming from the solvation shell. While this might limit the precision of the code in the case of solvated proteins, previous works (Kynde *et al.*, 2014; Skar-Gislinge *et al.*, 2015) suggest that in the case of an MP embedded in a nanodisc it doesn’t seem to provide a key contribution to the intensity and it would only add a seemingly unnecessary complication. Indeed, the model should feature already sufficient freedom in the number parameters even without the need to explicitly taking into account the solvation shell. In the case were further evidence might suggest a fundamental contribution coming from the solvation shell, this assumption can be easily dropped by introducing solvent beads (Svergun *et al.*, 2001). Many successful strategies to do this have been developed, exploiting different levels of coarse-graining of the water molecule representation and different methods to build the solvation shell around the protein (Svergun *et al.*, 1995; Park *et al.*, 2009; Grishaev *et al.*, 2010; Stovgaard *et al.*, 2010; Grudinin *et al.*, 2017).

As a final remark, we want to discuss what we consider to be the strongest assumption behind our approach, which is however shared by all the other ab-initio strategies. As known, the measured scattering intensity represents an average over an ensemble of, possibly very different, molecular configurations. Nonetheless, ab-initio methods try and fit the experimental signal with a single, say *average*, configuration. In the case of rigid molecules, this will not make a fundamental difference, as the protein conformational space will be, by definition, rather limited. For flexible proteins (e.g. tissue factor), however, this *average* configuration might not strictly represent a low energy conformer or even a member of the conformational ensemble at all and the aforementioned assumption might make the interpretation of the results particularly non-trivial. We advise the users to keep this in mind when using the *Marbles* software. A possible way to account for structural fluctuations at the level of ab-initio model would be, taking inspiration from applications of the Maximum Entropy principle to biomolecular systems (Pitera & Chodera, 2012; Cavalli *et al.*, 2013; Boomsma *et al.*, 2014), to implement a *multi-replica* version of the methods, where none of the replicas is forced to fit exactly the experimental signal but rather only their average computed intensity.

### 5.2. Comparison with existing methods

The literature is rich in techniques to retrieve the structure of soluble proteins from SAXS signals based on ab-initio bead modelling. The approach described in this work, as anticipated in section 5.1, shares many common features with GASBOR (Svergun *et al.*, 2001) and DAMMIN (Svergun, 1999), which were among the first algorithms to solve the problem, but extends their capabilities by including also an explicit treatment of a membrane nanodisc.

There are a few recent examples of alternative software approaches to solve the more general problem of retrieving an unknown membrane protein structure from SAS data. The MONSA program from the Svergun group (Svergun, 1999) was probably the first to demonstrate how an *ab initio* approach could be used to restore multiphase structures from different SAS contrasts, e.g. obtained through contrast variation SANS on protein DNA complexes. However, as discussed more recently (Koutsioubas, 2017; Molodenskiy *et al.*, 2020), the original unconstrained version of MONSA does not produce sufficiently uniform and reliable solutions in the case of embedded membrane proteins. Indeed we have come to the same conclusion when investigating a similar full bead-based approach to solve the problem for nanodisc embedded MPs (unpublished work).

The more recent software MEMPROT (Pérez & Koutsioubas, 2015) had the specific goal to model the component of the SAS signal provided by the detergent surrounding the protein. The original version of the code required that the high-resolution structure of the studied membrane protein was already known. But a new release, named DANVILLE (Koutsioubas, 2017), extended its capabilities by also modelling the protein structure likewise through a bead modelling approach when at least two scattering contrasts, e.g. two SANS curves at different solvent contrasts, were available. DANVILLE then co-refines the combined MP and detergent corona, assuming both are unknown.

Very recently, the Svergun group developed a pipeline for the fast analysis of detergent embedded membrane proteins on the fly during a SAXS beamtime. The pipeline searches for prior knowledge about the studied membrane protein: if its structure is already known, the data are analysed through MEMPROT as referred above. If other sufficiently strong prior knowledge is available, e.g. about scattering length densities of membrane protein and detergent headgroups and tails, then MONSA is applied in combination with these constraints along with constraints about thickness of the detergent corona, i.e. following a strategy resembling that described in the DANVILLE software. In the case where no prior knowledge is available, the pipeline simply resorts to using a combination of DAMMIF and DAMMIN on the low-q data to obtain an overall representation of the size and shape of the complex (Molodenskiy et al., 2020). While the membrane protein structures obtained through both the modified MONSA and the MEMPROT applications of the pipeline both look plausible (Molodenskiy *et al.*, 2020), the approach reported less convincing model fits to the (notoriously challenging) high-*q* part of the small-angle scattering data which are dominated by the scattering from the detergent corona.

Like the above discussed approaches, *Marbles* also relies on a combination of SAS data and prior knowledge. As discussed also in our previous articles (Kynde *et al.*, 2014; Skar-Gislinge *et al.*, 2015) we consider this a necessity when restoring MP structures from small-angle scattering data. With the structural homogeneity of the nanodiscs, our accurate structural description of the empty reference discs, and the plausible assumption that a single pass membrane protein will only provide a minor pertubation of the disc structure, we will argue that the approach employed in *Marbles* leaves us with less uncertainty with respect to separating out the protein contribution to the data. This should provide us with higher structural resolution and more reliable protein structures. We also note that we obtain a much better agreement between model and experimental data than the obe obtained by Molodenskiy and collaborators (Molodenskiy *et al.*, 2020). This however comes at the cost that the *Marbles* software is presently less general than the above approaches, in that it focuses only on single pass membrane proteins. Therefore, since *Marbles*, MEMPROT/DANVILLE and the membrane protein pipeline of the Svergun group are adapted to slightly different purposes, a direct comparison between them is not possible.

Besides bead modelling approaches, also other techniques have been applied to determine protein shapes. In particular, DENSS employs a model-free approach to retrieve a protein’s electron density, by iteratively modelling and anti-Fourier transforming 3D scattering intensities (Grant, 2018). Despite its predictive power in the case of solvated proteins, it remains uncertain whether the same approach is generally robust for membrane proteins or other multiphase systems. Indeed, even if the possibility to allow for negative contrasts is present in the code, to the extent of our knowledge no applications of this approach to membrane proteins have been published.

Given the similarities, but also the striking differences with previous works in the literature, we believe that *Marbles* is a useful alternative to determine the shape of a protein in a nanodisc.

### 5.3. Possible extensions

As a matter of principle, the open software philosophy behind *Marbles* allows for the implementation of case- or user-specific penalty functions. Beyond system specific modification, however, we foresee some particularly useful integrations that can be implemented into future releases of the code.

First of all, it would be useful to include further constraints through contrast variation SANS penalty terms. This would reduce the intrinsic SAS uniqueness problem by the introduction of further experimental information. The implementation of such terms is rather straightforward and we have indeed used the approach for similar problems in the past (Skar-Gislinge et al., 2010; Kynde et al., 2014), but the code has to be adapted accordingly and made efficient.

Secondary, as we anticipated in section 5.1, a possible extension of the code goes in the direction of implementing a replica bead modelling algorithm. The rationale behind this is the fact that a single structure doesn’t necessarily reflect the conformational ensemble contributing to the experimental SAXS intensity. Therefore, the code might be extended to allow for multiple replicas to run simultaneously and contribute to the 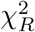 without being forced to fit the signal individually, but only on average. The penalty function acting on replica *r* would then take the form:

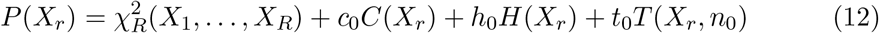

where *R* is the total number of replicas and the 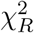 defines the discrepancy between the experimental intensity and the average intensity of the replicas.

Thirdly, a generalisation of *Marbles* to encompass other types and shapes of the carrier particles and in particular to various types of detergent micelles or polymer stabilised carriers, is desirable. This is also the target addressed in the approach described by Molodenskiy and collaborators (Molodenskiy *et al.*, 2020). Such a generalisation is feasible and in principle trivial from the modelling point-of-view. However, the fulfilment of the assumption that the carrier particle should be relatively monodisperse and have the same conformation with and without the embedded membrane proteins unfortunately limits the broader practical relevance of this path to a small set of carrier systems.

Finally, we want to mention the possibility to modify the code engine. While simulated annealing is a robust method which has been implemented with great success in commonly used softwares (Svergun, 1999; Svergun *et al.*, 2001), the computational efficiency of the code and the robustness in the determination of the minimum of the penalty function could be increased by making the code compatible with existing powerful minimization routines, e.g. maximum-likelihood based Monte Carlo (Ferkinghoff-Borg, 2002) and Bayesian generalized ensemble Markov chain Monte Carlo (Frellsen *et al.*, 2016).

## 6. Conclusions

In this manuscript we introduced a new software, *Marbles*, capable of modelling the shape of a membrane protein embedded in a nanodisc through the mere knowledge of the corresponding SAXS intensity, the protein FASTA sequence and a SAXS based model of the unloaded nanodisc particle. We discussed the range of applicability of the code and compared it with existing studies in the literature, concluding that, despite its intrinsic limitations, *Marbles* has a wide range of applicability and provides new features that are lacking in existing bead modelling softwares. We tested the performance of the code on toy models and on two relevant examples, consisting of two proteins with very different topologies. The obtained results, in both cases compatible with the current experimental and computational knowledge on the systems under investigation, show that *Marbles* is a reliable software for shape determination and that it is able to distinguish among different topologies without having to manually adjust the running parameters.

## Appendix A Generation of synthetic data

To build the synthetic SAXS data mentioned in section 3.2, we adopted the following strategy. First, we fitted the experimental SAXS intensity of a solution of empty nanodiscs (discussed in section 4.1) using *WillItFit* in order to deduce realistic fit parameters for the analytical nanodisc model. Secondarily, we used the PDB of cytochrome P450 3A4 (CYP3A4) built by Denisov et al. (Denisov et al., 2012) as our reference protein. Using this PDB and the analytical model we defined two scenarios: (i) one where the protein is placed in the center of the nanodisc (model 1); (ii) one where the protein is in the proximity of the nanodisc’s rim (model 2). Model 1 was built by displacing the protein in such a way that the intramembrane helix was completely immersed in the nanodisc while the center of mass of the protein’s extracellular domain was found approximately in *x* = 0 and *y* = 0, protruding from the axis provided by the normal to the nanodisc plane. Model 2 was built similarly, but the protein was shifted towards the border of the lipid bilayer. We then employed *WillItFit* to obtain the SAXS intensities of the two models. In order to consider realistic errors and noise into the data, we associated to each intensity point the experimental error *δI* obtained by measuring the SAXS intensity of a solution of nanodisc bound CPY3A4 proteins (see section 4.1). After that, we resampled each intensity point assuming Gaussian statistics within the errorbars: 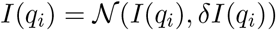, where 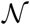 is the normal distribution, *I*(*q_i_*) represents its average and *δI*(*q_i_*) its standard deviation.

## Appendix B Toy Models

In this section we will go in detail in explaining how our strategy to determine the initial position of the random sphere of beads makes Marbles minimization more efficient. To show that the procedure is able to determine a reasonable starting position on the nanodisc, we employed the two synthetic SAXS intensities of cytochrome P450 3A4 discussed in appendix A. We initialized Marbles with a random sphere of beads placed in (*x* = 0, *y* = *z*_0_) and moved it by fine grained steps of 1 Å in *x* and *y* in order to cover all the rectangle surrounding the nanodisc. For each point on the grid we computed the corresponding value of 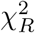. In both the cases of model 1 and 2, the initial optimization was able to determine an initial position which is very close to the exact one (see Fig. 5 (a) and (b), right column).

**Fig. 5.**
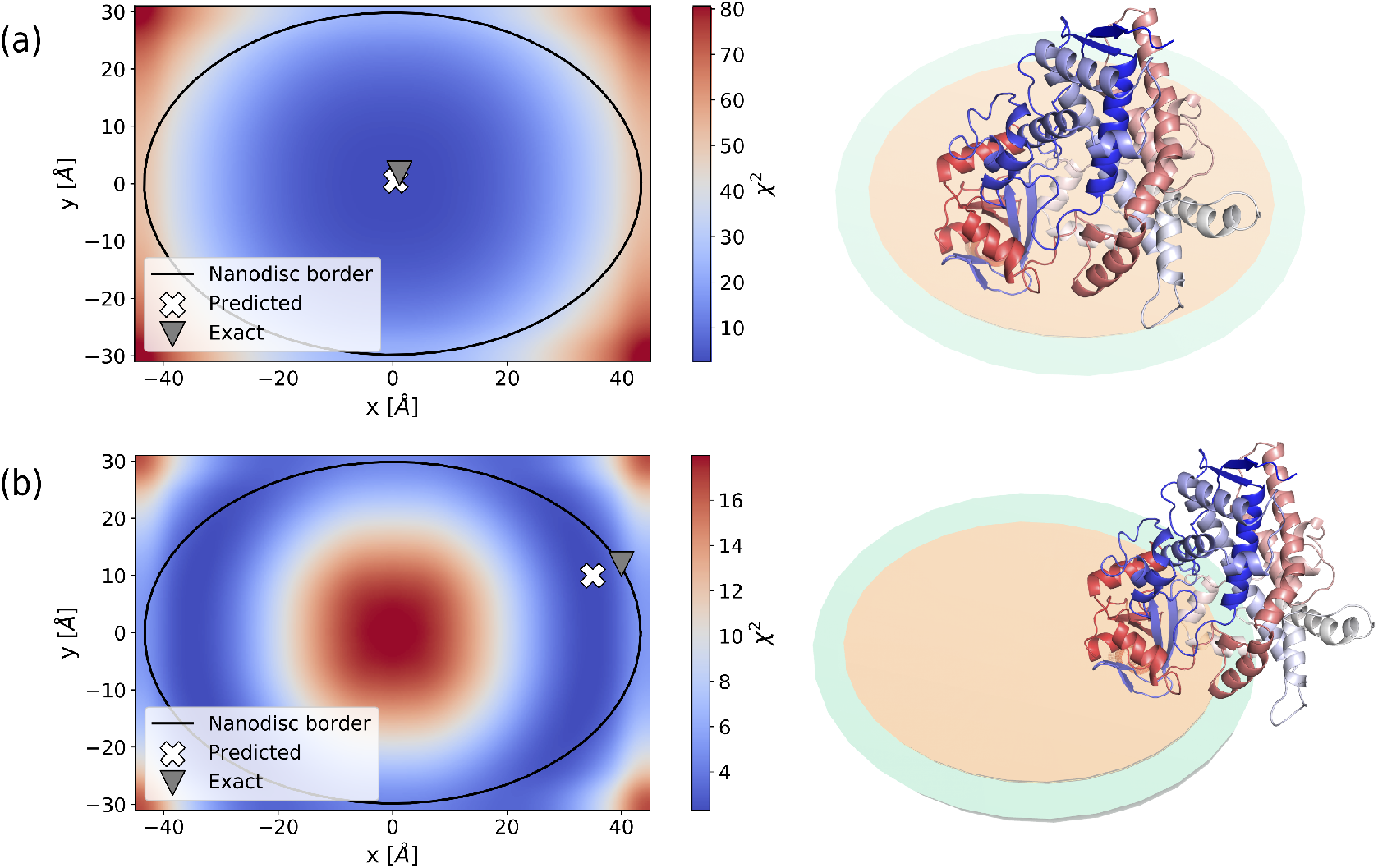
Validation of the center of mass (COM) optimization strategy. Each point on the heatmaps corresponds to the center of mass coordinates of the randomically generated sphere of beads, while the colors are employed to show the 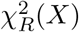 computed between the scattering intensity of the sphere-plus-nanodisc system and and synthetically generated SAXS data. The black solid line encloses the lipid bilayer surface as seen from the top, the grey triangle represents the COM’s location of the target protein, while the white cross highlights the predicted minimum of the 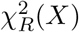. In (a) the reference protein is placed in the center of the nanodisc (see the protein’s cartoon representation on the right). In (b), the reference protein is instead placed on the nanodisc’s rim (see the protein’s cartoon representation on the right).

To test the robustness of this estimation and assess how much it is conserved throughout the whole optimization run, we performed four *Marbles* runs using both models 1 and 2. In all the runs we employed the parameters shown in Tab. 1. In Fig. 6 we report a summary of the results. In the case of model 1 (Fig. 6 (a)), the center of mass of the final models is not located exactly in correspondence of (*x* = 0, *y* = 0), but remains nonetheless confined in the wide, low 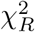 region in the middle of the nanodisc. The average distance of the results from the nanodisc center, and therefore from the center of mass of the target protein, is about 15Å.

**Fig. 6.**
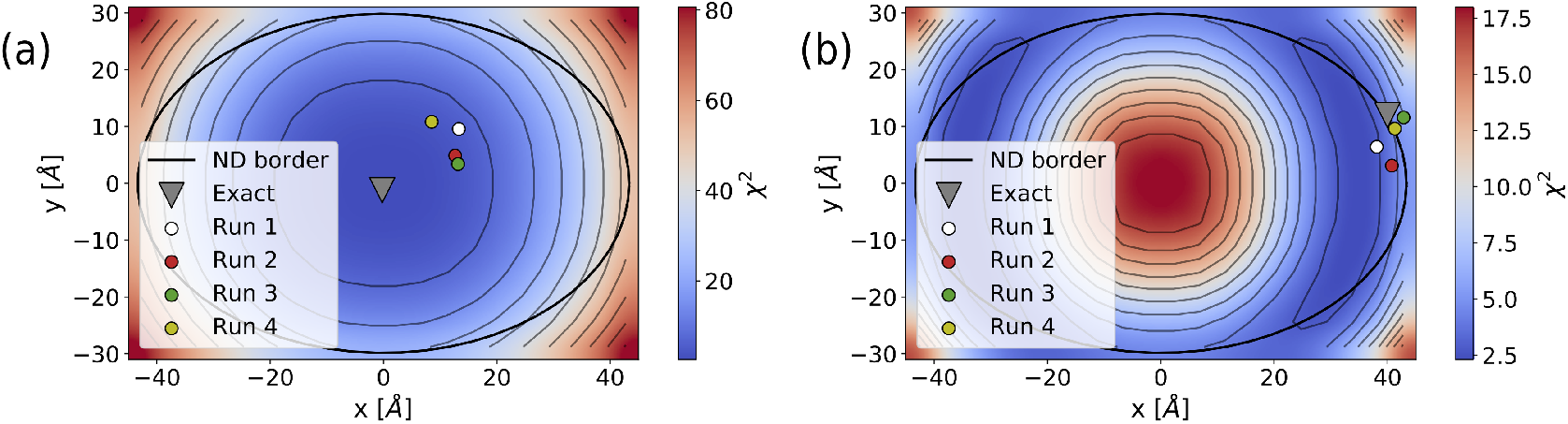
Validation of the center of mass (COM) optimization strategy. The heatmap shows the 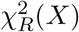 landscape obtained from the center of mass optimization of the initial random sphere (see section 3.2 and Fig. 5 for further details) in the case of (a) model 1 and (b) model 2. The black solid line encloses the lipid bilayer surface as seen from the top and the grey thin lines show the level curves of the heatmap. The grey triangle represents the COM’s location of the target protein, while the colored circles show the final location of the COM of four *Marbles* runs. In both (a) and (b), the heatmap represents only the results obtained for Run 1, but the overall features for the other runs remain identical.

While this might seem like a substantial shift, we have to ask ourselves whether such a precise information about the protein’s location can be encoded in the orientationally averaged SAXS intensity. In particular, most of this information is found in the maximum diameter of the system *D*_max_. We expect that, in a case like this where the nanodisc is larger than the membrane protein, a shift of the order of 10Å from the center of the nanodisc should not substantially alter the value of *D*_max_. To confirm this, we performed an analysis of the scattering density functions *p*(*r*). To obtain them, we first aligned the all-atom structure of CYP3A4 to the optimal models obtained with *Marbles* using DAMAVER. In this way, we obtained four different constructs with centers of mass shifted by approximately 15Å from the nanodisc center. We then applied the same methodology described in appendix A to compute four synthetic SAXS intensities. We finally employed BayesApp (Hansen, 2012) to compute the corresponding *p*(*r*) functions (Fig. 7), and compared them with the densities obtained for the target constructs of model 1 and 2. It is clear that *D*_max_ between the four runs and the target reference of model 1 are absolutely compatible, while the *D*_max_ obtained for the target protein of model 2 is approximately 1 nm larger. This observation is compatible with our assumption that it is hard to detect the exact location of the protein when it is placed centrally on the nanodisc. This is also underlined by the slowly decreasing values of the 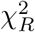 on the nanodisc’s surface (Fig. 6 (a)) that, compatibly with our *D*_max_ analysis, provides a high level of ambiguity for the determination of the optimal position.

**Fig. 7.**
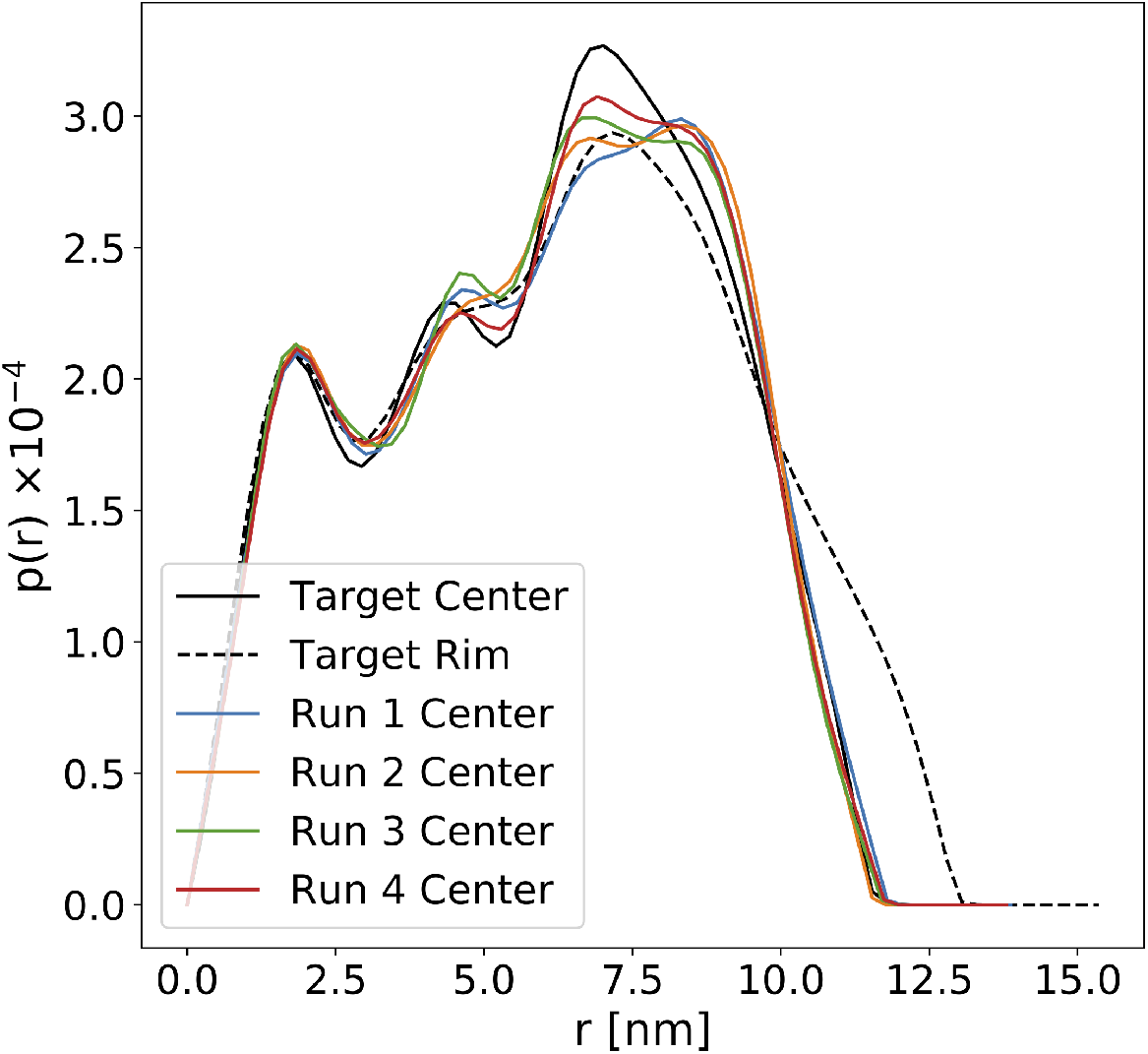
Colored solid lines: *p*(*r*) functions computed from the four different constructs with centers of mass shifted by approximately 15Å from the nanodisc’s center; black solid line: *p*(*r*) function computed from the target construct employed to define model 1; black dashed line: *p*(*r*) function computed from the target construct employed to define model 2.

In model 2 (Fig. 6 (b)), instead, the four runs find almost the exact location of the target protein. Indeed, the higher *D*_max_ in this case encodes the fact that larger structures need to be built to fit the data, and *Marbles* is able to correctly interpret this information by placing the protein close to the nanodisc’s rim.

Finally, for sake of completeness we report in Fig. 8 the typical results obtained by *Marbles* applied on the synthetic data representing model 1 and 2. In both cases, Marbles is able to predict a protein shape compatible with the one of the target protein that was used to generate the data.

**Fig. 8.**
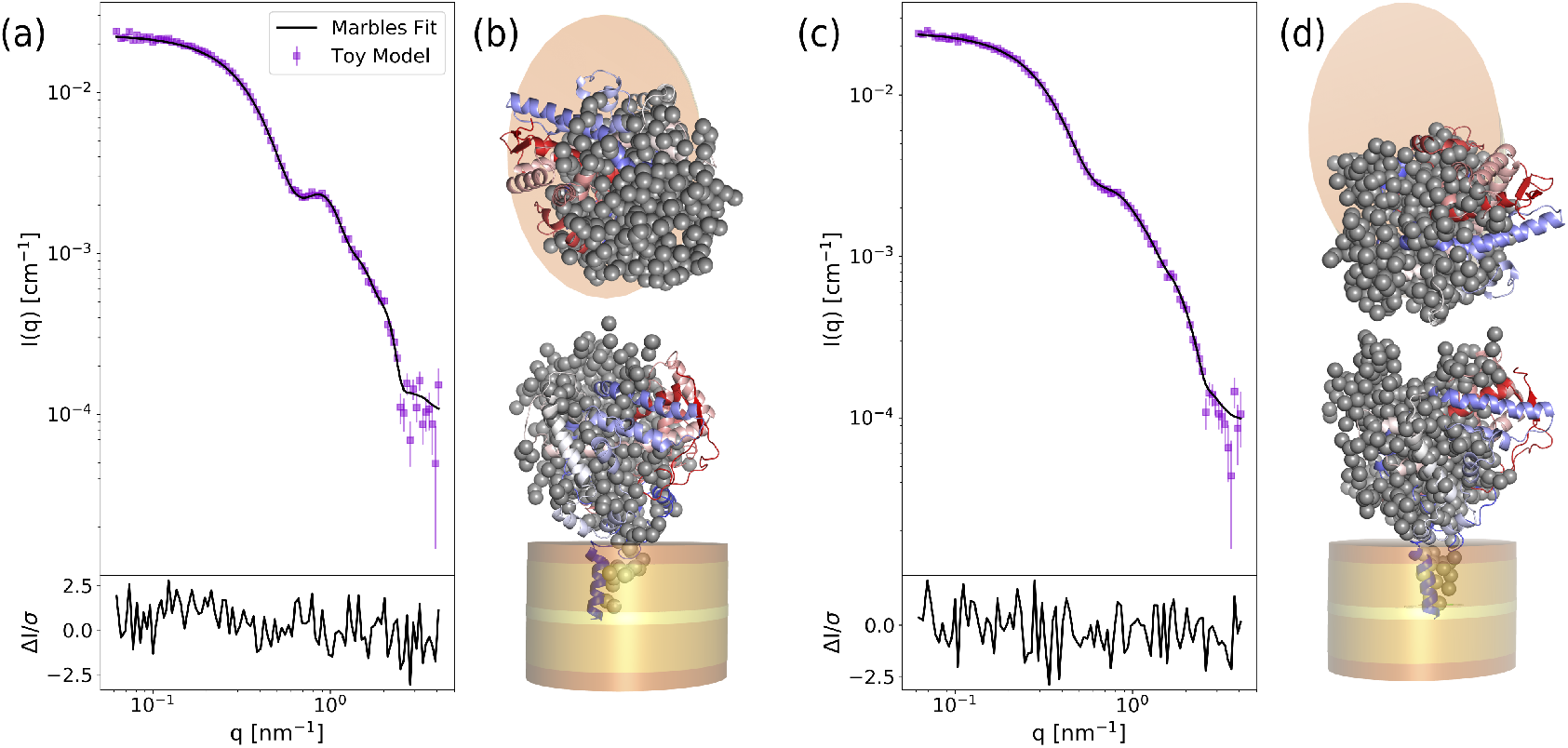
Results of the application of *Marbles* to the two synthetic models. SAXS curves are shown as purple dots in (a) for model 1 and in (b) for model 2. representing the synthetic models (a) 1 and (c) 2. In both cases, the full black curves show the corresponding model fits. The lower panels report the residuals between the synthetic and the computed signals. Typical configurations obtained by *Marbles* as seen from the top and from the front are reported in (b) for model 1 and (d) for model 2. The PDB of the target protein, colored by residue index, is shown aligned to *Marbles* models.

## Appendix C Comparison with previous code

To assess the computational speedup obtained by *Marbles* with respect to the old version of the code (Skar-Gislinge et al., 2015), we tested both of them on the experimental data of CYP3A4 on a 2.3 GHz Intel Core i5 processor. On average over 10 runs, the original code required approximately 50 minutes to converge, while *Marbles* took less than 5, meaning that we were able to achieve approximately a 10-fold speedup. This speedup comes both from an optimization of the source code and from the initial optimization of the guess protein configuration. Indeed the original code, where this strategy is not implemented and the initial configuration is always a sphere located in (*x* = 0, *y* = 0), requires approximately 80 passes to converge, *Marbles* always requires only 60. This means that most of the initial high-temperature stage of simulated annealing in the original code is spent diffusing over the nanodisc’s surface, as clearly shown in Fig. 9.

**Fig. 9.**
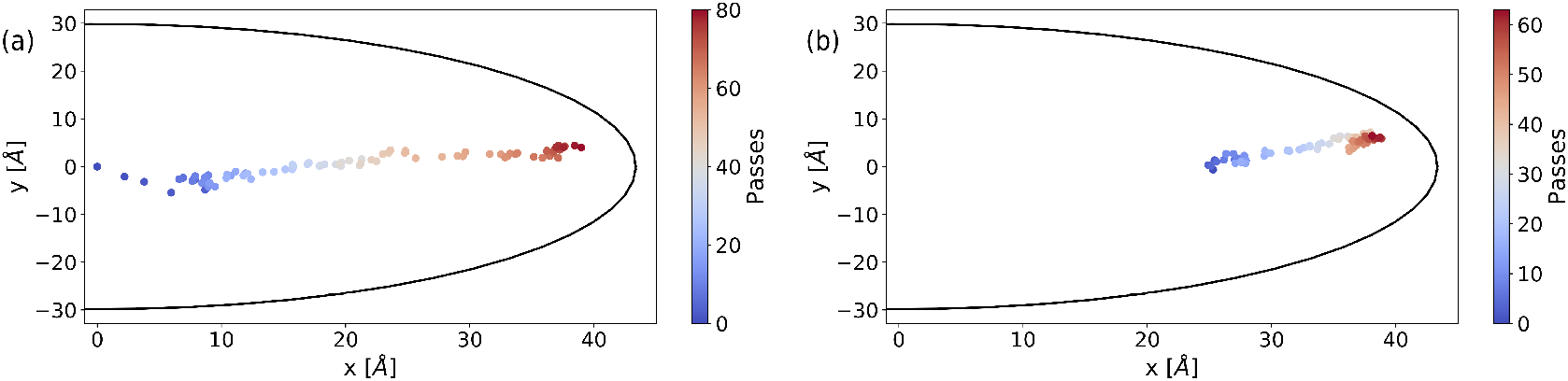
Example of the displacement of the center of mass of the protein during a single run of simulated annealing in the case of (a) the original code (Skar-Gislinge *et al.*, 2015) and (b) *Marbles.* The black solid line shows the left nanodisc border while the colormap is used to represent the number of simulated passes.

## Appendix D Input, parameters and output

The mandatory ingredients required by *Marbles* are: (i) the protein’s sequence, in FASTA format; (ii) the SAXS intensity of the protein embedded in the nanodisc; (iii) the SAXS intensity of a solution of empty nanodiscs; (iv) a *WillItFit* file with the results of the fit of the empty nanodisc system; (v) an estimate of the number of residues composing the intramembrane portion of the protein *n*_0_. While still necessary, the SAXS intensity of a solution of the empty nanodisc will not be employed by *Marbles* directly, but rather to *WillItFit* to determine the best nanodisc parameters. Other inputs can be provided but are not mandatory, such as the coupling constants *c*_0_, *h*_0_, *t*_0_ in Eq. 11, the cooling schedule *s* and the starting and convergence temperatures 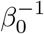 and 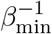. The default settings of such parameters, however, should perform well for the majority of the systems (see Tables 1 and 2 for the numerical values and appendix E for a discussion on how default parameters have been selected).

The output of the code consists in several files: (i) the protein configurations at the end of each pass, in PDB format; (ii) the computed SAXS intensities at the end of each pass; (iii) a log file which summarizes the parameters employed in the run; (iv) a summary of the values of the penalty functions throughout the whole run. While most of the output files are mostly used for diagnostics (and the output can be therefore suppressed), the last intensity file provides the best fit of the model and the last PDB structure represents the best 3D protein shape recovered by *Marbles*.

## Appendix E Choice of parameters

The default settings for *c*_0_, *h*_0_ and *t*_0_ parameters have been selected in such a way that the initial phase of the penalty function optimization is dominated by the minimization of *C*(*X*), *H*(*X*) and *T*(*X*) (approximately the first 10 passes, as shown in Fig. 10). This ensures that the protein will remain compact and connected while the fine details of the SAXS intensity are optimized during the second phase of the run. The relative strength of these three penalties are set in a such a way to make them approximately of the same order of magnitude throughout the whole minimization run.

**Fig. 10.**
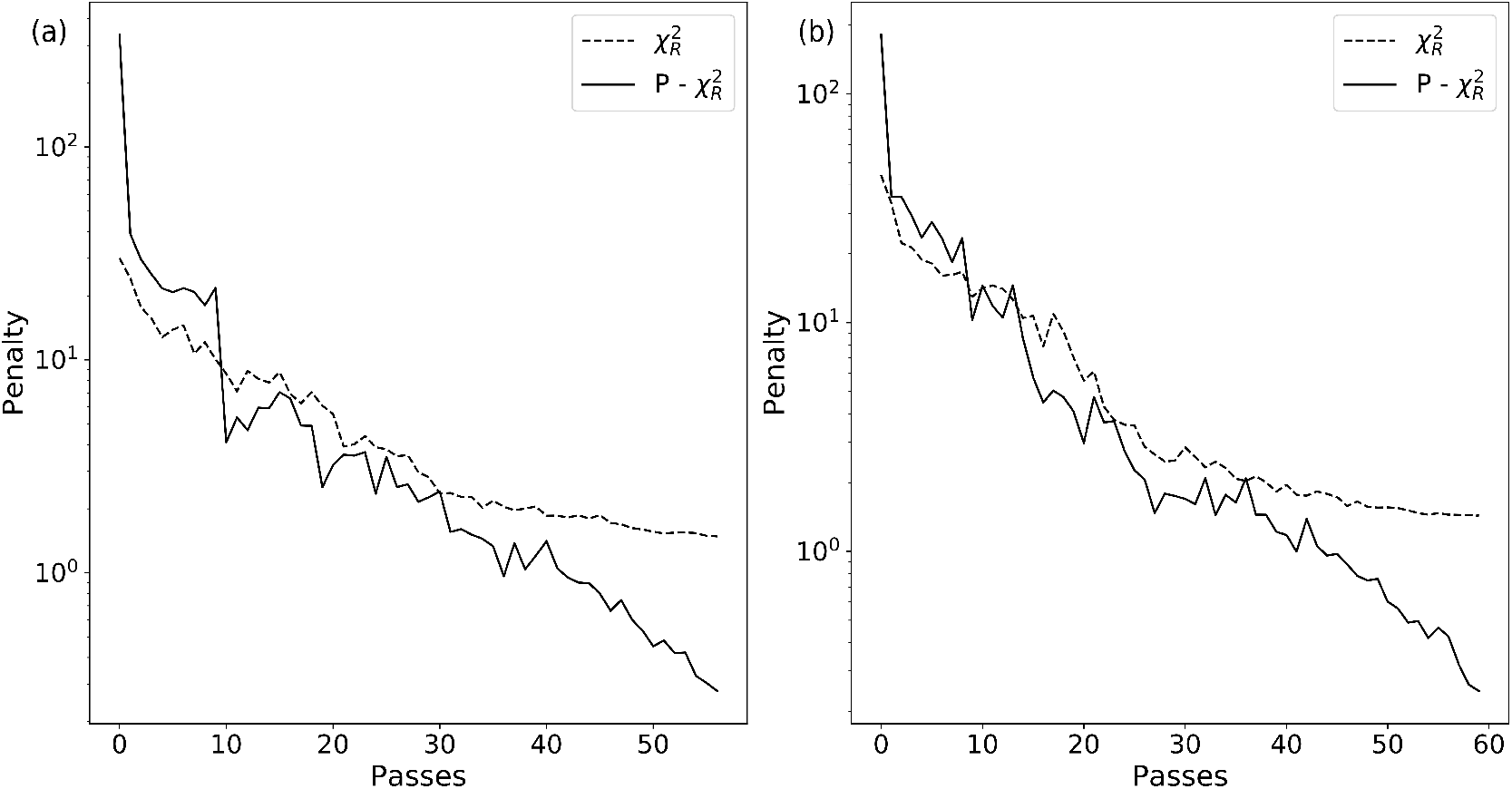
Behavior of the penalty functions in the case of (a) synthetic model 2 and (b) application to CPY3A4.

Concerning the effective temperature, we set the initial one to 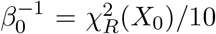, while the cooling schedule s is set to 0.9. While s = 0.9 is a rather standard option for simulated annealing (Svergun *et al.*, 2001), the choice of 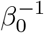 is less trivial and in our case has been made empirically. Nonetheless, this choice has the advantage of being automatically system dependent, so that the initial effective temperature doesn’t need to be manually adjusted when changing from system to system. We also tested different options for the determination of 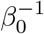 and for the cooling schedule, none of which have provided better results or appreciably faster convergence in our tests (see e.g. sections 2.1 and 2.2 in Ref. (Park & Kim, 1998)). We do not exclude, however, that other and even better choices of the parameters might be possible.

We thank Martin Nors Pedersen, Raul Araya Secchi and Nicolai Johansen for inspiring discussions; Martin Cramer Pedersen for discussions and useful suggestions on the manuscript; Nicholas Skar-Gislinge for providing SAXS data for Cytochrome P450, the version of the code used in the original publication and support on it; and finally Frederik Grønbek Tidemand for providing SEC-SAXS data for Tissue Factor.

## Synopsis

*Marbles* is a novel software that employs SAXS intensities to predict the shape of membrane proteins embedded into membrane nanodiscs.

